# Hep3D: A 3D single-cell digital atlas of the liver to study spatio-temporal tissue architecture

**DOI:** 10.1101/2023.01.21.525037

**Authors:** Dilan Martínez, Valentina Maldonado, Cristian Pérez, Rodrigo Yañez, Valeria Candia, Yannis Kalaidzidis, Marino Zerial, Hernán Morales-Navarrete, Fabián Segovia-Miranda

## Abstract

Three dimensional (3D) geometrical models are not only a powerful tool for quantitatively characterizing complex tissues but also useful for probing structure-function relationships in a tissue. However, these models are generally incomplete due to experimental limitations in acquiring multiple (>4) fluorescent channels simultaneously. Indeed, predictive geometrical and functional models of the liver have been restricted to few tissue and cellular components, excluding important cellular populations such as hepatic stellate cells (HSCs) and Kupffer cells (KCs). Here, we performed deep-tissue immunostaining, multiphoton microscopy, deeplearning techniques, and 3D image processing to computationally expand the number of simultaneously reconstructed tissue structures. We then generated a spatio-temporal singlecell atlas of hepatic architecture (Hep3D), including all main tissue and cellular components at different stages of post-natal development in mice. We used Hep3D to quantitatively study 1) hepatic morphodynamics from early post-natal development to adulthood, and 2) the structural role of KCs in the murine liver homeostasis. In addition to a complete description of bile canaliculi and sinusoidal network remodeling, our analysis uncovered unexpected spatiotemporal patterns of non-parenchymal cells and hepatocytes differing in size, number of nuclei, and DNA content. Surprisingly, we found that the specific depletion of KCs alters the number and morphology of the HSCs. These findings reveal novel characteristics of liver heterogeneity and have important implications for both the structural organization of liver tissue and its function. Our next-gen 3D single-cell atlas is a powerful tool to understand liver tissue architecture, under both physiological and pathological conditions.

## Introduction

The liver fulfils a wide range of functions, including metabolism, detoxification, protein synthesis, and production of biochemicals that aid digestion. These diverse functions rely on an intricate 3D tissue architecture where different cell types coexist and interact in a coordinated fashion, and thus, triangulating the exact spatial location of a cell is key to understanding its role. Macroscopically, the liver is composed of functional anatomical units called lobules, which include cellular and tissue structures located between the central vein (CV) and the portal triad (hepatic artery, portal vein (PV), and bile duct). The blood coming from the gut, pancreas, and spleen enters the liver via the PV and mixes with blood from the hepatic artery. Blood then flows toward the CV through a highly branched network of blood vessels called sinusoids, whereas, bile flows through a second bile canaliculi (BC) network in an antiparallel direction. The space between the central and the portal veins is filled predominantly by hepatocytes and non-parenchymal cells. Hepatocytes constitute the primary cell type at the core of liver function and are responsible for processing blood and secreting bile into the BC. They are “sandwiched” between the sinusoidal endothelial cells and share the apical surface with multiple neighboring hepatocytes to form a 3D BC network. This organization allows the hepatocytes to have numerous contacts with the sinusoid and BC networks to maximize the exchange of molecules(1). Non-parenchymal cells also play important roles in liver function and growth regulation. They include liver sinusoidal endothelial cells, hepatic stellate cells (HSC), and Kupffer cells (KC). HSCs are located in the space of Disse between sinusoids and hepatocytes, store Vitamin A, and secrete most of the extracellular matrix(2). Indeed, activation of HSCs is a central driver of several liver diseases(2). KCs are liver specific self-renewing resident macrophages located inside the sinusoidal capillary that play an important role in initiating hepatic immune responses and clearing circulating endotoxins(3). Increasing evidence suggests that hepatocytes, HSCs, and endothelial cells are in close contact, thereby, forming the so-called ‘hepatic niche’ (4,5). Therefore, it is not surprising that changes in the function of any cell integrated within the niche could eventually impact their neighborhood.

Classical histology has played a crucial role in understanding liver tissue structure. It is simple, versatile, and extremely accessible. However, this technique also presents several disadvantages, 1) it is not quantitative, 2) overlooks 3D information, 3) poorly distinguishes non-parenchymal cells, and 4) some tissue structures are not visible e.g., the BC network. Developments in tissue clearing, high-resolution fluorescence microscopy, and 3D image analysis have allowed 3D liver tissue reconstruction in the form of geometrical models (6,7), to describe liver tissue architecture with unprecedented detail. Over the last few years, geometrical models have proven to be a game-changer and have illuminated basic principles of liver tissue organization. Some of the main findings include: i) hepatocytes display a pronounced spatial zonation based on their ploidy (6), ii) the first predictive model of biliary fluid dynamics(8), iii) hepatocytes polarity exhibit liquid-crystal order(9), iv) liver regeneration after partial liver hepatectomy requires biomechanical growth control(10), and v) 3D reconstruction of human liver biopsies from non-alcoholic fatty liver disease patients show profound topological defects in the 3D BC network that lead to zonated micro-cholestasis(11).

Till date, fluorescence microscopy is usually limited to a maximum of four fluorescent markers. Due to this restriction, 3D geometrical models have not been able to describe all the main cell types and tissue structures simultaneously, thus overlooking the organization of the hepatic niche. Simultaneous reconstruction of all important tissue networks (BC and sinusoids), cellular (hepatocytes, HSCs, and KCs) and sub-cellular components (nuclei) would require at least six different markers, making it impossible to observe all the structures of interest at once. Here, we combined deep tissue immunostaining, optical clearing, multiphoton microscopy, deep learning techniques, and 3D image processing to virtually expand the number of markers and generate a spatio-temporal 3D single-cell atlas of the liver tissue, Hep3D. We used Hep3D to describe morphological changes established during early postnatal development and the structural role of KCs in liver tissue architecture. This atlas provides a powerful tool to quantitatively describe each cell type, its spatial organization, and its possible cross-interactions. Hep3D will help to identify (sub)structural characteristics of liver architecture, providing a quantitative tool to understand both liver biology and pathology with unprecedented detail.

## Results

### SeeDB optical clearing shows high compatibility with different staining modalities while preserving tissue morphology

To generate a 3D geometrical tissue model containing all the main cell types and tissue components, we first optimized our pipeline of deep tissue imaging. Our standard pipeline involved several steps including, fixation, vibratome sectioning, staining, optical clearing, imaging at high resolution using multiphoton microscopy, and 3D reconstruction with the software Motion Tracking(6). For staining specific structures, we used a CD13 antibody for BC, Flk-1 antibody for sinusoids, and the small molecule dyes, DAPI (4,6-diamidino-2-phenylindole) and phalloidin, for nuclei and cell borders (actin mesh), respectively. There are several optical clearing techniques, however, many of them are compatible with only a subset of markers or change the tissue morphology (e.g. tissue expansion)(12). Moreover, as most of these techniques were developed for the brain, there is scant information about their use in liver tissue(13). To find the optical clearing method that best suited our requirements, liver slices were stained and optically cleared using different methods including, FOCM(14), FRUIT(15), RTF(16), SeeDB(17) and SeeDB2G(18). We focused on well-stablished clearing methods that have been shown to preserve tissue morphology and tested them side-by-side in terms of staining compatibility and preservation of tissue morphology. Most of the clearing methods showed high compatibility with antibody staining, while their performance was variable with the small molecule dyes (Supplementary figure 1a). Even though the methods evaluated here showed different optical characteristics (Supplementary figure 1b), no major differences in tissue transparency were appreciable when 100 μm liver sections were compared (Supplementary figure 1a). It is likely that differences in transparency may be appreciable in thicker samples. Finally, we compared the effect of the optical clearing on tissue morphology both macroscopically (e.g. liver slice expansion) and microscopically (e.g. BC radius) (Supplementary figure 1c,d). FRUIT was the only method that resulted in tissue expansion both macro and microscopically. We found that SeeDB was the best clearing method for 100 μm tissue slices, showing high compatibility with different types of staining while maintaining tissue morphology.

### Virtual tissue labeling enables simultaneous 3D reconstruction of liver tissue components

Fluorescence microscopy is usually limited to 4–5 markers to avoid bleed-through of the fluorescence emission. We overcome these experimental constraints by generating virtual 3D images of the BC and sinusoidal networks based on the phalloidin staining using deep convolutional neural networks (CNNs) (Figure 1a-c and Supplementary figure 2). We then used the remaining channels to visualize KCs and HSCs, using the markers F4/80 and desmin, respectively (Figure 1b,c). We expanded our 2D CNN-based toolbox for the prediction of BC and sinusoids from phalloidin-stained images (19) to a fully 3D model. Our method showed remarkable accuracy when comparing the virtual with the real BC and sinusoidal networks (Supplementary figure 2a-d). The virtual BC and sinusoidal networks showed a high signal-to-background ratio (Supplementary figure 2b,d) and their morphometric properties were very similar to the original networks (Supplementary figure 2e,f). This approach allowed us to image and reconstruct all the main components of the liver tissue microarchitecture simultaneously, thus generating a multi-parametric 3D single-cell atlas of the liver which we named Hep3D (Figure 1d and Supplementary movie 1).

**Fig. 1:**
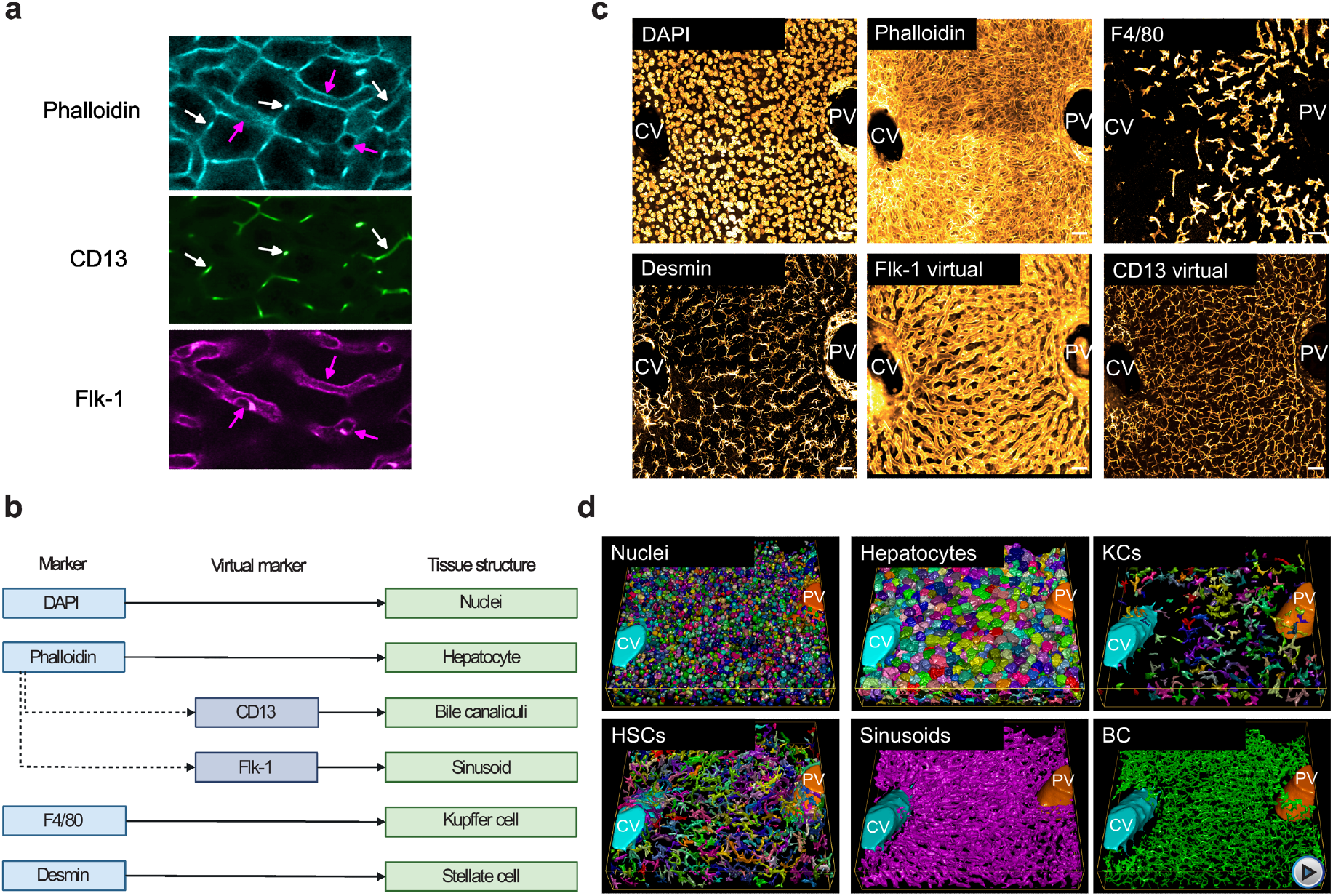
Deep tissue imaging and convolutional network allow complete reconstruction of liver tissue structure. (a) Images from 100 μm liver section stained with Phalloidin (cell border), Flk-1 (sinusoids) and CD13 (bile canaliculi). White and magenta arrows indicate the BC and sinusoidal networks respectively. (b) Real and virtual markers used for each cell type and structure. (c) Maximum projection of a 30 μm z-stack covering an entire CV-PV axis. Scale bar 30 μm. (d) 3D reconstruction of the main structures of the liver tissue. Central vein (light blue), portal vein (orange), nuclei (random colors), hepatocytes (random colors), KCs (random colors), HSCs (random colors), sinusoids (magenta) and bile canaliculus (green).

### 3D single-cell atlas reveals morpho-spatial differences in liver tissue architecture from early post-natal development to adulthood

Next, we determined morphological changes in liver tissue from postnatal day 1 (P1), postnatal day 16 (P16), and adult mice. These temporal stages were selected as they encompass key phases in early postnatal development during which the liver undergoes several morphological changes. In P1 mice, the hepatocytes are mostly diploid and remnants of embryonic development such as hematopoietic cells can be detected(20,21). In P16 mice, the liver tissue architecture was partially mature, as mice were weaned, triggering an increase in hepatocyte ploidy. Adult mice displayed a mature and well-established liver tissue architecture. Quantification of several morphological features including tissue (BC and sinusoidal networks) and cellular (hepatocytes, KCs, and HSCs) components is summarized in Supplementary tables 1–3.

Both BC and sinusoidal networks were fully connected at all stages. Whereas BC radius decreased, the sinusoidal radius increased over time. The BC network corresponds to ~5% of tissue volume at all ages analyzed, however, the sinusoidal network increased from 14% in P1 to 25% in adults (Supplementary table 1). Hepatocytes occupied around 58% of the tissue volume in all mouse stages analyzed. Their morphology was homogenous in P1 and gradually became more heterogenous in terms of nuclearity, ploidy, and cell volume as the mice aged. Hepatocyte cell volume changed from 2152 ± 160 μm^3^ in P1 mice to 4470 ± 525 μm^3^ in adults. The increase in cell volume correlated with an increase in the number of polyploid cells (41% in P1, 48% in P16 and 73% in adults). Other morphological properties, such as the percentage net contribution of apical, lateral and basal membranes, and the number of neighbors, showed only modest changes (Supplementary table 02). HSCs were highly elongated at all ages groups investigated. While the volume of individual HSCs increased with age, the nuclear volume decreased. Surprisingly, the number of HSCs reduced by ~42% from P1 to adult while they occupied a relatively constant proportion of the tissue (7-9%) (Supplementary table 3). F4/80 positive cells were very elongated, and their number reduced as the mice aged (99307 cells/mm^3^ in P1, 25810 cells/mm^3^ in P16 and 13345 cells/mm^3^ in adults). We also detected a significant reduction in the volume occupied by KCs during neonatal development (8.75% in P1, 6.95% in P16 and 3.57 % in adults) (Supplementary table 3). It is probable that a large fraction of the F4/80 positive cells in P1 represents not only KCs but also macrophages located within erythroblastic islets, which disappear from the liver about one week after birth in mice(22,23).

Even though analyzing the cells in the liver as a population provided an extremely informative overview of the tissue, a detailed description of liver tissue structure has to take into account possible changes in morphology along the CV–PV axis(6,8,24,25). Therefore, we computationally divided the CV–PV axis into ten equidistant regions and quantified the different morphological properties within each sub-region. While the spatial distribution of the sinusoidal radius appears homogeneous at all stages, the BC showed a modest increase towards the veins only in adults (Figure 2a). Mono-nucleated diploid (1×2n) and bi-nucleated tetraploid (2×2n) hepatocytes were enriched toward the CV and PV, while hepatocytes with higher ploidies tend to be concentrated in the middle zone of adult livers (Figure 2b), in agreement with previous reports (6,26). Our data suggested that the spatial arrangement of hepatocytes according to their ploidy is an event that occurs after weaning (Figure 2b). The spatial distribution of KCs and HSCs showed a clear anti-correlated pattern along the liver lobule for all ages. While KCs were enriched in the middle zone, HSCs were concentrated around the big veins (Figure 2c). Our data supports previous findings where it has been shown that the average values, although important, hide important aspects of tissue architecture.

**Fig. 2:**
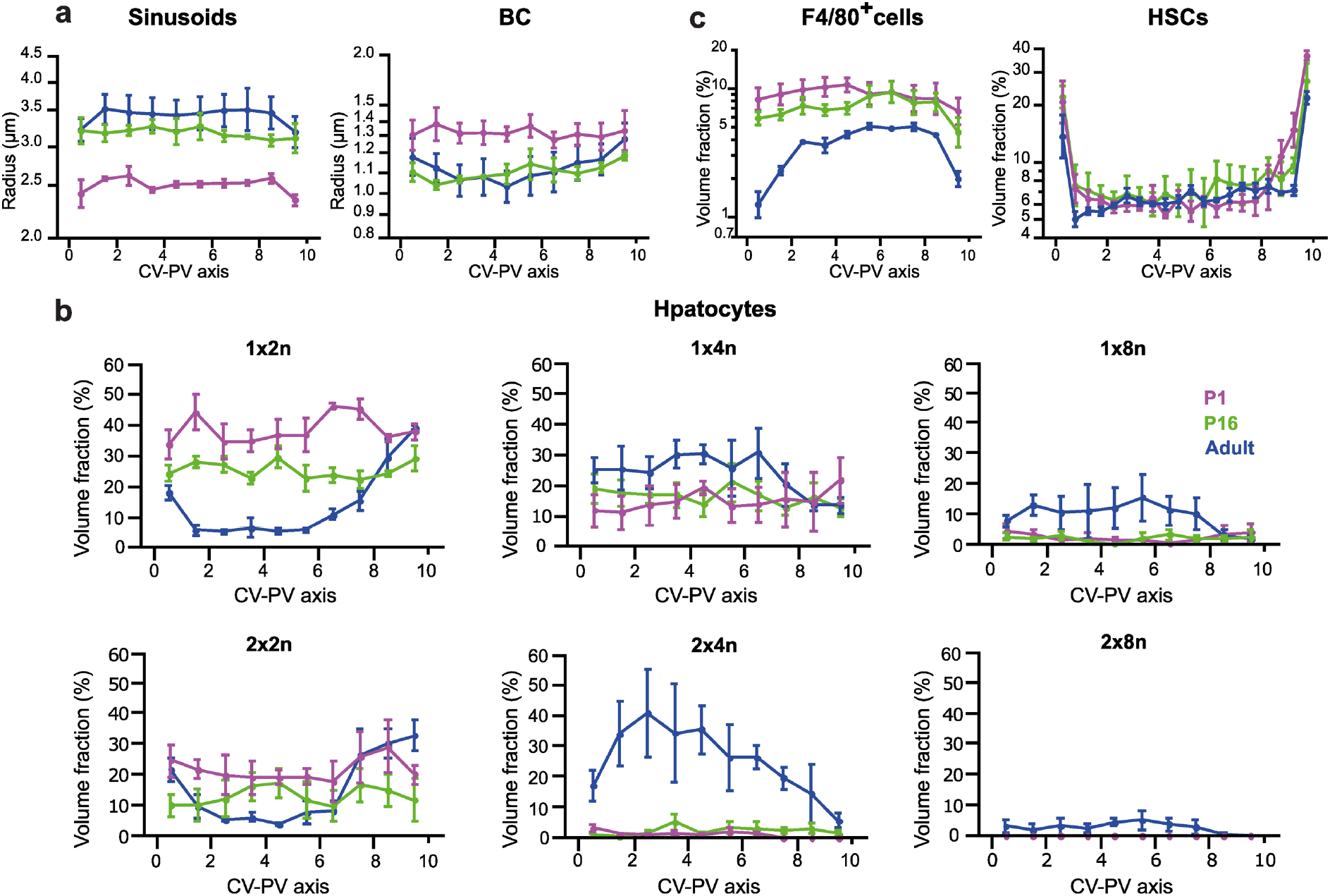
Morphological changes occurring during early postnatal development revealed by spatial analysis. (a) Spatial distribution of sinusoid and BC networks radius. (b) Quantification of the percentage of tissue volume occupied by hepatocytes with different ploidy along the CV–PV axis. (c) Quantification of the percentage of tissue volume occupied by F4/80+ cells and HSCs. In (a) and (c) the “y” axes are in log-scale. The measurements were taken from the CV to the PV, subdividing the space into 10 equal parts. P1 = 3 samples, P16 = 3 samples, Adults = 3 samples. Quantification represented by mean ± s.e.m.

### Hep3D unveils unexpected spatial interaction between HSCs and KCs

Having different cell types and tissue components simultaneously in our 3D reconstruction allowed us to do a systems level analysis of hepatic structure. First, we computed the percentage of hepatocyte surface in contact with other tissue structures. While some areas of contact remained constant as the mice aged (HSCs, BC), the contact with other tissue components either increased (sinusoids) or decreased (neighboring hepatocytes, KCs) (Figure 3a). Next, we explored the possibility of physical contact between HSCs and KCs and observed that they were in close proximity (Figure 3b). We quantitatively corroborated this observation by measuring the number of cell-cell contact sites between the HSCs and KCs. We found that each KC, on average, had 2–3 contact sites with HSCs. The number of contact sites did not change as the mice aged (Figure 3c). We then wondered if the contacts were made via the long protrusions emanating from HSCs or the body of the cells (i.e., close to their nuclei). To address this question, we measured the distance between nuclei of different cell types (Figure 3d). Analysis of inter-nuclear distance distribution showed that the vast majority of KC nuclei were remarkably close to HSC nuclei (Figure 3d). Indeed, we found that a large fraction (40 ± 5%) of KCs have nuclei close to the HSC nuclei (inter-nuclear distance smaller than 2 μm) (Figure 3e). This is not the case for hepatocyte nuclei, which are separated by at least 7–8 μm, on average, from the HSC and KC nuclei. These results imply that the proximity of HSC-KC nuclei is non-random, suggesting the existence of a strong direct interaction between KCs and HSCs.

**Fig. 3:**
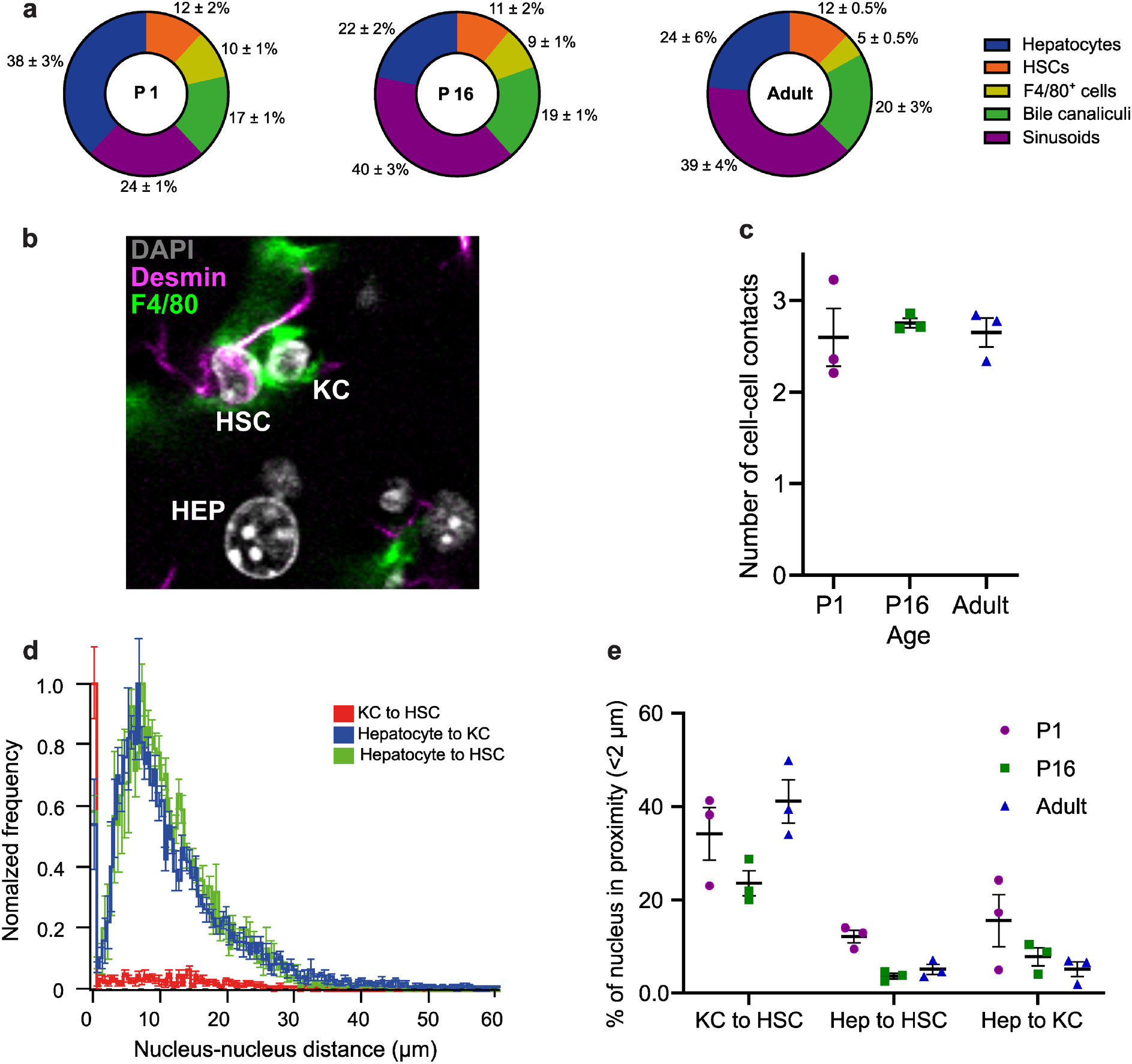
Interplay between cell types and tissue components. (a) Surface percentage of the hepatocytes in contact with the different structures (BC and sinusoids) and cells (HSCs, F4/80+ cells and other hepatocytes) present in the liver. (b) A maximum projection of 30 μm from the tissue showing the staining of desmin (red), F4/80 (green), Flk-1 (blue) and DAPI (magenta). (c) Average number of KC contact sites on the HSCs. (d) Normalized distribution of nucleus-nucleus distance between hepatocytes, KCs and HSCs. (e) Percentage of nuclei from different cell types which are closer than 2μm. P1 = 3 samples, P16 = 3 samples, Adults = 3 samples. Quantification represented by mean ± s.e.m.

### KC depletion alters the liver tissue architecture

To test if the physical proximity and contact between cells are linked to functional cell-cell interactions, we ablated the KC population and evaluated the effects of this perturbation on the general tissue architecture and the spatio-temporal organization of all cell types. Briefly, KCs were depleted from the liver by intravenous injection of liposome-encapsulated clodronate(27,28). To allow the KC-depleted liver tissue enough time to develop hepatic tissue architecture, while causing minimal distress to the mice, we first estimated the frequency of injections. Mice were retro-orbitally injected with clodronate and tissue samples were analyzed on days 1, 3, 5 and 7 post-injection (Supplementary figure 3). We observed that KCs started repopulating the liver on day 7, and therefore we injected the mice with clodronate every 5 days for long-term depletion experiments. Next, mice received one injection every 5 days from P16 to P30 (Figure 4a-b). Clodronate treatment achieved a 91% reduction in the number of KCs/mm^2^ (Figure 4c-d). At the structural level, the absence of KCs for 15 days (P16 to P30) did not cause detectable alterations on the BC and sinusoidal networks (Supplementary table 04). Even though we only performed a short-term depletion, it was enough to observe morphological changes in hepatocytes and HSCs. In the case of hepatocytes, most of their characteristics were unaffected except nuclearity and ploidy, both of which showed an increase upon KC depletion (Supplementary table 05 and figure 4e-g). This effect may be attributed to the previously described role of KCs on hepatocyte proliferation(29,30) and the correlation between cell division and hepatocyte ploidy(31). Interestingly, the tissue volume occupied by HSCs increased dramatically upon KC depletion (Figure 4h and Supplementary table 06). This was observed along the entire liver lobule (Figure 4h) and was accompanied by a massive increase in the number of HSC (Figure 4i). The HSC shape was also affected (Supplementary table 06). The number and shape of HSCs may indicate HSC activation in the absence of KCs (32). In concordance with previous reports, our morphological analysis of liver tissue suggests the existence of a direct crosstalk between KCs and HSCs (33,34).

**Fig. 4:**
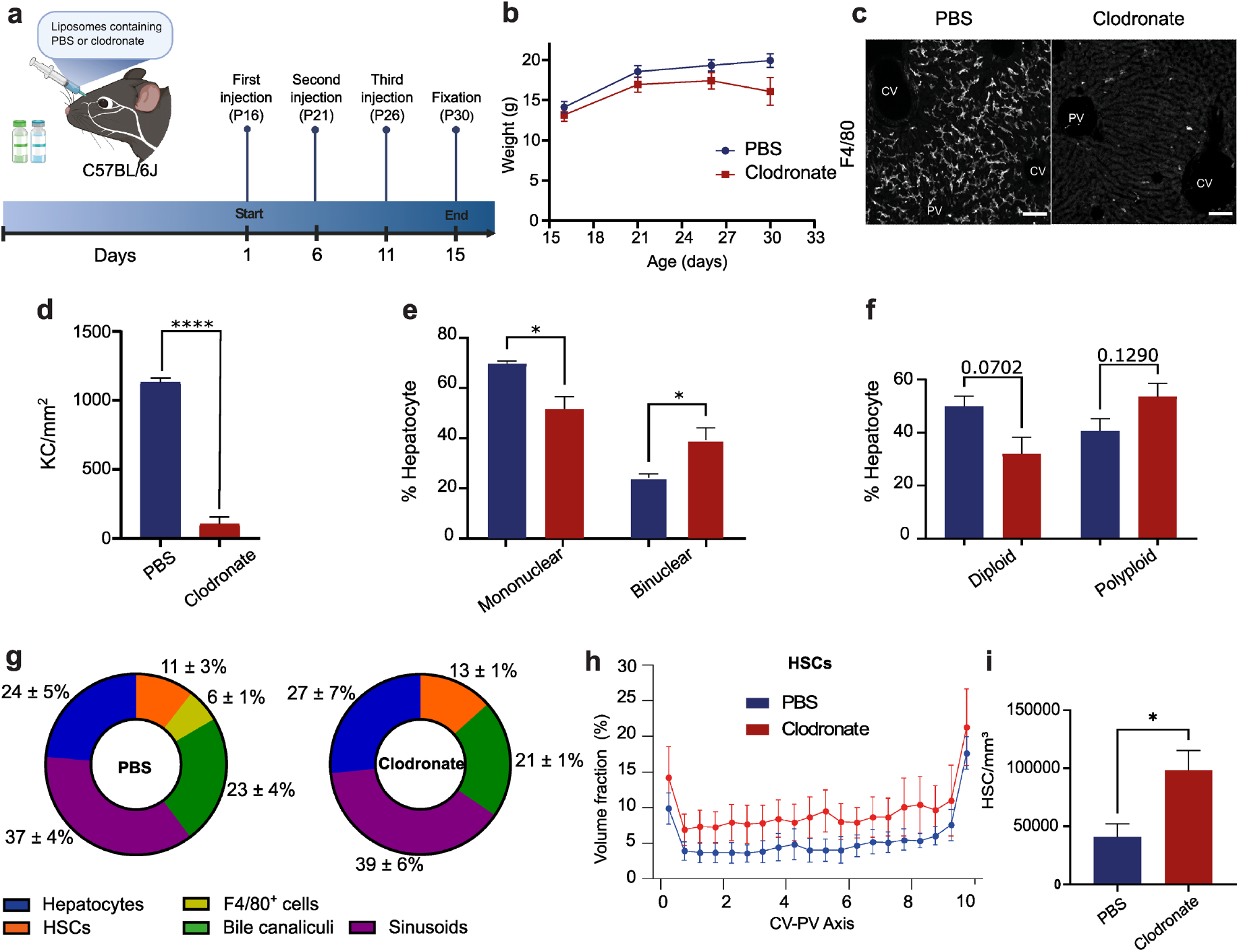
Structural defects of liver tissue architecture upon KC depletion. (a) Scheme showing the KC depletion experiment. Mice were injected with liposomes containing clodronate or PBS (control), every 5 days, starting at P16 until P30. (b) Mice were weighted every 3 days from the start to the end of the experiment. (c-d) 100 μm slices were stained with anti F4/80 and KC were quantified. Percentage of (e) mononuclear and binuclear, (f) diploid and polyploid and hepatocytes in PBS and clodronate conditions. (g) Percentage of hepatocyte surface contacting with other cell types and tissue structures. (h) Spatial distribution of HSCs volume fraction from PBS and clodronate. (i) Comparison of total number of HSCs/mm^3^ in PBS and clodronate. PBS = 3 samples, Clodronate = 3 samples. Quantification represented by mean ± s.e.m. Two-tailed t-test (*P<0.05, **P<0.01, ***P<0.001, ****P<0.0001).

## Discussion

Geometrical models have proven to be powerful and versatile tools to quantitatively describe the morphology of liver micro-architecture (BC and sinusoids)(8,35,36) and its cells (hepatocytes, KCs, and HSCs)(6,37). Unfortunately, it has not been possible to image and reconstruct all these structures simultaneously due to technological limitations. In this study, we showed that a combination of deep tissue imaging, deep learning and traditional 3D image analysis techniques enables the 3D reconstruction of the main structural components of liver tissue architecture simultaneously. Our 3D single-cell digital atlas of the liver single-cell, Hep3D, includes hepatocytes, HSCs, KCs (both nuclei and cell surfaces), BC, and sinusoidal networks. Hep3D enabled us to extract the spatio-temporal evolution of morphological tissue characteristics during different post-natal stages. Since different metabolic and morphological features change along the developing liver lobule(6,8,24,25), our 3D atlas allows a deeper understanding of the spatially heterogeneous phenomena (e.g. liver zonation). As a proof of concept, we described how liver tissue structure changes during the physiological transition from post-natal early development to adulthood and the structural role of KCs in this process. In concordance with previous reports, we found that hepatocyte ploidy increased as the mice aged and occupied different regions of the liver lobule based on this characteristic(26,38,39). We also observed an unexpected anti-correlated spatial distribution of HSCs and KCs. The crosstalk between these cells became even more evident following KCs ablation from the liver which had a profound impact on the number and morphology of the HSCs. Hep3D provides a holistic yet in-depth overview of liver tissue organization, with the potential for detecting even subtle changes in liver microarchitecture.

The combination of experimental data with computational models of tissues has proven successful in revealing how different tissues and organs function(8,40). Volumetric imaging and optical clearing techniques play crucial roles in this process(41). In this scenario, tissue preservation is paramount, as data extracted from the 3D reconstructions can not only be used to get a quantitative understanding of tissue morphology but also, as an input for mathematical models. In this study, we evaluated different optical clearing techniques to achieve volumetric imaging and tissue preservation. SeeDB showed the highest compatibility with our staining method and the lowest impact on tissue morphology. Indeed, 3D reconstruction of BC showed morphometric parameters indistinguishable from samples without optical clearing. Unfortunately, SeeDB is not the best technique in terms of transparency so if a specific application requires the user to image deeper into the tissue, the development of new methodologies or the use of new microscopy techniques (i.e. 2-photon and light-sheet microscopy) will be necessary(41).

Another important component of the pipeline is accurate 3D reconstruction of the tissue. Even though they are a powerful and versatile tool, their efficient generation still poses some difficulties. The main challenges that we faced were related to the accurate segmentation of densely packed irregular shaped nuclei, and the segmentation of cells with complex shapes e.g. HSCs. The field of artificial intelligence is revolutionizing bioimage analysis and is actively providing new tools to overcome these problems(42). In the future, it will be interesting to explore methods such as Stardist to improve our 3D reconstructions (43), Cellpose(44), Plantseg(45) etc.

Recently, a lot of effort has been devoted to understanding tissues at the cellular level(46), even integrating spatial information(47), e.g. single-cell spatial transcriptomics. While these technologies are a breakthrough in cell biology at the tissue scale, they usually overlook tissue morphology and niche organization, and consequently, their role in tissue biology and pathology. One example is our finding about the spatially anti-correlated organization of the KC and HSCs, wherein regions with a high density of HSCs showed a lack of KCs, and conversely in the regions with low density of HSC, KCs are more abundant and located in close proximity of the HSCs. Strikingly, we observed a clear pattern of spatial distribution of the nuclei of these cells in 40% of the cases, where HSC and KC nuclei are within close proximity (less than 2 μm). The biological implications of this observation are unknown, showcasing the potential of Hep3D in the generation of biological questions based on morphological observations. We expect that changes in the functional activity of the cells constituting the hepatic niche, could eventually impact the other cell members. Considering the crucial role that KCs and HSCs play in liver diseases (48,49), Hep3D will provide a key tool to survey the principles of liver tissue organization under diverse conditions.

This versatile tool also has vast potential for advancing our understanding of liver diseases. A previous version of our 3D reconstruction, which did not include non-parenchymal cells, helped discover a set of cellular and tissue parameters correlated with non-alcoholic liver disease progression. Moreover, we previously 1) discovered profound defects in the BC network that only became apparent by 3D analysis, and 2) integrated the morphometric analysis of the BC network with mathematical models to create personalized biliary fluid dynamic simulations(11). In this context, a multiparametric image that includes a quantitative tissue perspective and the crosstalk with key cellular constituents represents a promising future technology to study liver pathology i.e. progression from steatosis to hepatocellular carcinoma. Hep3D is a valuable and versatile tool for future investigations on the quantitative description of liver tissue microarchitecture in a wide variety of physiological and pathological conditions.

## Methods

### Animals

Postnatal day 1, 16, and adult C57BL/6J mice were obtained from the animal facility (Centro Regional de Estudios Avanzados para la Vida (CREAV)) at the Universidad de Concepción. The animals were maintained in strict pathogen-free conditions and received *ad libitum* feeding. All procedures performed were approved by the vice rectory of ethics and biosecurity committee from the investigation and development of Universidad de Concepción.

### Sample collection and immunostaining

Mice livers were fixed through intracardiac perfusion with 4% paraformaldehyde 0.1% Tween-20/PBS and post-fixed overnight with the same solution at room temperature. In the case of P1 mice, the livers were collected and fixed by immersion in 4% paraformaldehyde 0.1% Tween-20/PBS over 5 days at room temperature. 100 μm thick liver sections were obtained with a vibratome. Immunolabeling (Supplementary table 07) and optical clearing were performed as described previously (9).

### Evaluation of optical clearing methods

Once the immunolabeling was performed, liver tissue sections were cleared by different methods. The protocols used included SeeDB2G(18), SeeDB(9,17), FRUIT{Hou.2015} and FOCM{Zhu.2019}. Samples in PBS (uncleared) were used as a control. To estimate the macroscopic changes in the size of the tissue, the overall area was measured by drawing the contour of the liver slice before and after the clearing using the software Fiji(50). Additionally, to measure microscopic changes, BC were stained and segmented and their radius was quantified using the software Motion Tracking(6). The difference in both areas and the BC radius were compared to the control condition to determine if the tissue shrunk or expanded.

### Imaging

Liver samples were imaged (0.3 μm voxel size) in an inverted multiphoton laser-scanning microscope (Zeiss LSM 780) using a 40×1.2 numerical aperture multi-Immersion objective (Zeiss). DAPI was excited at 780 nm using a Chameleon Ti-Sapphire 2-photon laser. Alexa Fluor 488, 555 and 647 were excited with 488, 561 and 633 laser lines and detected with Gallium arsenide phosphide (GaAsp) detectors.

### Image processing

The different components of liver tissues (BC, sinusoids, nuclei, HSCs, KCs and Hepatocytes) were reconstructed from high-resolution (voxel size 0.3 × 0.3 × 0.3 μm) fluorescent image stacks (≈ 100μm depth). To cover the entire CV-PV axes, 2×1 tiles were stitched using the image stitching plug-in of Fiji(51). All images were reconstructed using the software Motion Tracking (http://motiontracking.mpi-cbg.de) as described in (6). Briefly, for the pre-processing of the 3D images were first denoised using the PURE-LET algorithm(52) with the maximum number of cycles. Then, a background and shading correction was performed using the tool BaSiC(53) along the stack. Finally, all channels were aligned to a reference one using the function Correct 3D Drift from Fiji. To generate virtual images of BC and sinusoidal networks from, the images of phalloidin, we used a 3D extension of the deep convolutional neural network toolbox described in (19). The structures were segmented using a maximum entropy local thresholding algorithm. Artifacts generated by the segmentation (holes and tiny isolated objects) were removed by standard morphological operations (opening/closing). The triangulation mesh of the segmented surfaces was generated by the cube marching algorithm and tuned using an active mesh approach. To separate nuclei we used interactive watershed plug-in in Fiji(51) and a splitting algorithm in Motion Tracking(6). In the case of tubular structures, representations of the “medial axis” or “skeleton”, also called central lines, of the networks were generated. Central lines were generated as 3D graphs. HSCs and KCs were reconstructed based on the desmin and F4/80 staining. Finally, the shape of the cell surface (hepatocytes) was determined using an active mesh expansion from the reconstructed nuclei. For details, refer to (6).

### Morphological spatial analysis of cells and networks (BC and sinusoids)

The cell and nucleus elongation were defined as one minus the ratio of the mayor to minor elongation axis of the 3D object

Elongation = 1 - (A1/A3), where A1 and A3 correspond to the length of the maximum and minimum elongation axis of the 3D object

For details about other morphological quantifications, refer to (6,11).

### Clodronate treatment

KC depletion was achieved by macrophagic suicide(54,55). Briefly, 100 μl of clodronate liposome suspension (LIPOSOMA, CP-010-010, 5mg/ml) per 10 gr of the animal was retro-orbitally injected(56). As a control, mice were injected with a solution of liposomes and PBS. For long-term depletion experiments, mice were injected every 5 days.

### Statistical analysis

Data were analyzed using Prism 9 software.

## List of Abbreviations

CV: central vein
PV: portal vein
BC: bile canaliculi
HSC: hepatic stellate cells
KC: Kupffer cells
DAPI: 4,6-diamidino-2-phenylindole
CNN: convolutional neural network
P1: postnatal day 1
P16: postnatal day 16
P30: postnatal day 30

## Acknowledgments

We are grateful to Nirmalya Basu for a critical reading of the manuscript. We thank Constanza Aguirre for providing figures from Biorender. We would like to thank the following Facilities from Universidad de Concepción for their support: Centro de Microscopía Avanzada (CMA BIO-BIO) and Centro Regional de Estudios por la Vida (CREAV).

## Notes

**Financial support and sponsorship** This work was financially supported by Fondecyt (grant # 1200965).

**Conflict of interest** Authors declare no conflict of interest.

### Competing Interest Statement

The authors have declared no competing interest.

