## Supplemental material for "Hep3D: A 3D single-cell digital atlas of the liver to study spatio-temporal tissue architecture"

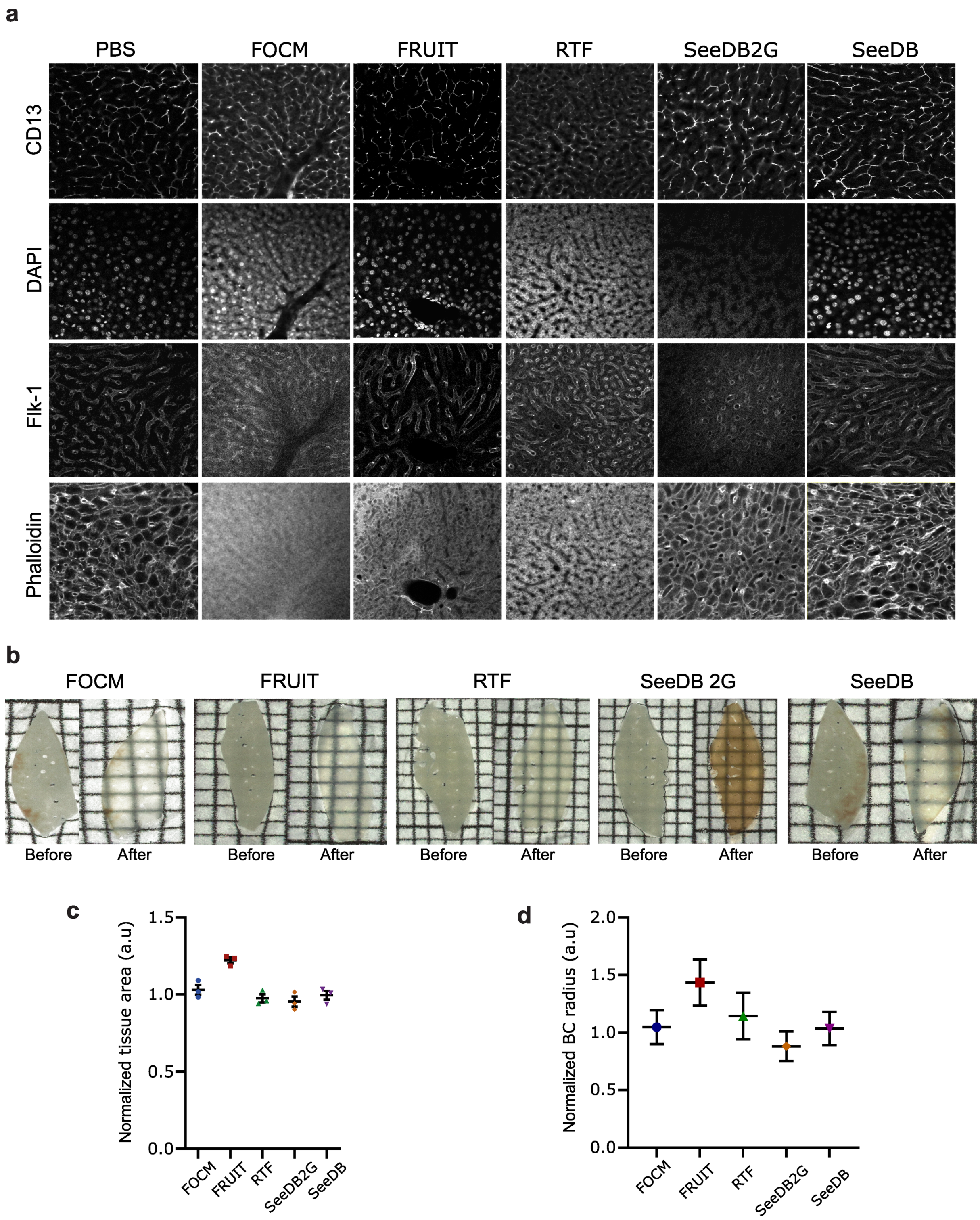


**Supp Fig. 1: Comparison of different optical clearing methods.**

(a) 100µm liver slices were stained with antibodies against CD13 and Flk-1, and the dyes phalloidin and DAPI. Different optical clearings methods were applied and the compatibility with the different markers was qualitatively evaluated. (b) Liver slices before and after the clearing are shown. (c) The outline of the liver slices was drawn in Fiji, the area was calculated and used as a readout of macroscopic tissue deformation. (d) BC was reconstructed with Motion Tracking and the radius was measured. The data were normalized to the PBS condition.

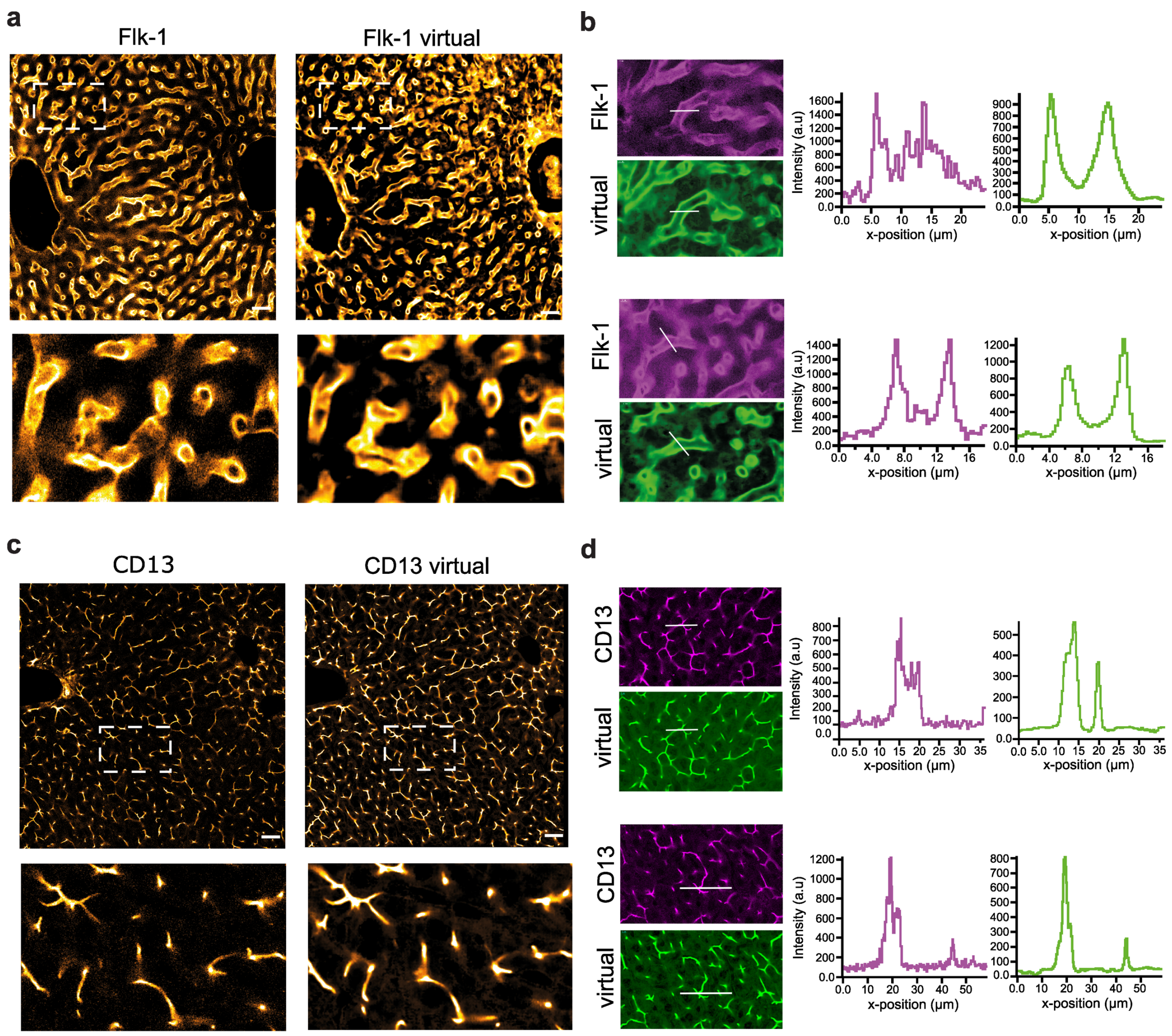


e

| Tissue feature | Sinusoids | Virtual sinusoids |
|---|---|---|
| Total length (m/mm3) | 4.26 | 3.44 |
| Volume fraction (%) | 16.8 | 16.7 |
| Radius (µm) | 2.9 ± 0.88 | 3.2 ± 1.08 |
| Conectivity (fraction) | 0.988 | 0.967 |
| Branch length (µm) | 15.9 ± 7.9 | 15.3 ± 8.01 |
| Junction density (number/µm) | 0.03 | 0.03 |

f

| Tissue feature | BC network | Virtual BC |
|---|---|---|
| Total length (m/mm3) | 6.5 | 6.0 |
| Volume fraction (%) | 3.46 | 4.5 |
| Radius (µm) | 0.91 ± 0.32 | 0.98 ± 0.42 |
| Conectivity (fraction) | 0.97 | 0.97 |
| Branch length (µm) | 10.3 ± 4.38 | 8.4 ± 4.96 |
| Junction density (number/µm) | 0.06 | 0.06 |

**Supp Fig. 2: Artificial neural networks can create images that resemble real staining.** (a) Representative images of the sinusoids from fixed mouse liver tissue. On the left, liver section stained with Flk-1. On the right, virtual image created with the 3D CNN-based toolbox from the phalloidin staining. Inset showing a magnification of the region highlighted on the upper image. (b) Intensity profiles along the lines drawn on two examples of real marker (magenta) versus virtual (green). (c) Representative images of the BC from fixed mouse liver tissue. On the left, liver section stained with CD13. On the right, virtual image created with the 3D CNN-based toolbox from the phalloidin staining. Inset showing a magnification of the region highlighted on the upper image. (d) Intensity profiles along the lines drawn on two examples of real marker (magenta) versus virtual (green). (e-f) Real and virtual images were 3D reconstructed and some morphometric properties were quantified.

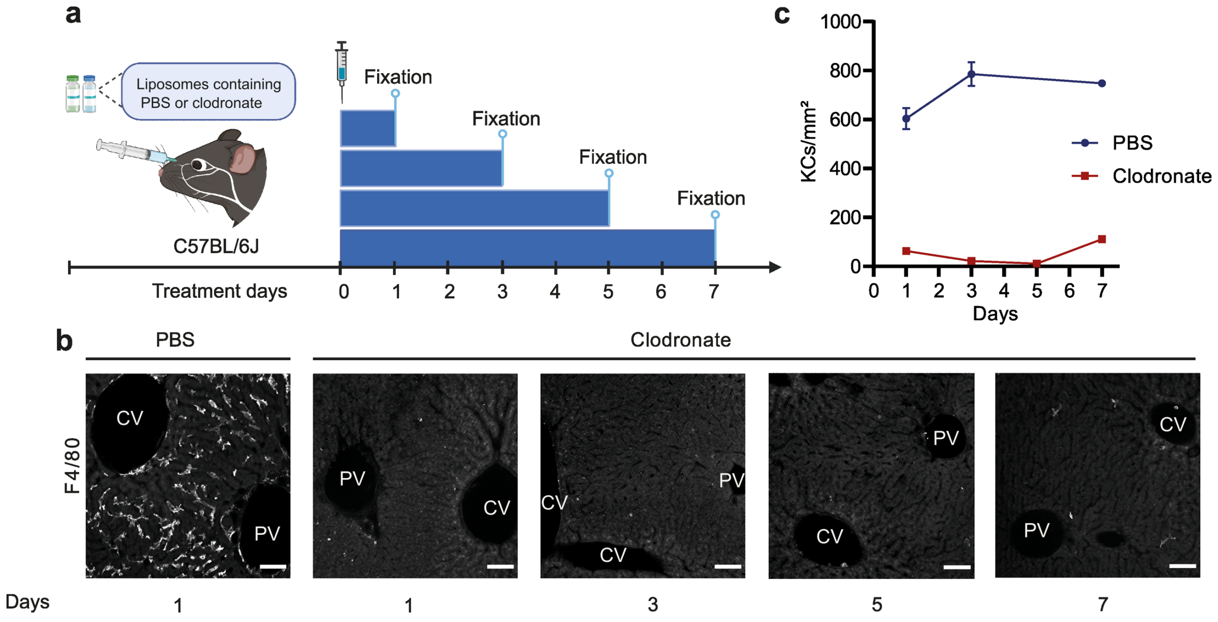


**Supp Fig. 3: Standardization of Kupffer cells depletion with clodronate liposomes.**
(a) Scheme summarizing the depletion experiment. A single clodronate injection was performed and liver samples were obtained after 1, 3, 5 and 7 days. (b) 100 µm thick liver sections were stained with anti-F4/80 antibody. (c) Number of KCs were quantified in Fiji using the images shown in (b) and were compared to control mice injected with PBS. PBS = 3 samples, Clodronate = 3 samples. Quantification represented by mean ± s.e.m.

**Supp movie 1: 3D single-cell morphometric atlas of liver tissue architecture.**
Central vein (light blue), portal vein (orange), sinusoids (magenta), bile canaliculus (green), nuclei (random colours), hepatocytes (random colours), HSCs (random colours) and KCs (random colours).

### Supp table 01: Morphometric analysis of BC and sinusoidal networks from early postnatal development to adulthood

| | Bile canaliculi | | | Sinusoids | | |
|---|---|---|---|---|---|---|
| | P1 | P16 | Adult | P1 | P16 | Adult |
| **Total length (m/mm³)** | 4.46 ± 0.5 | 6.17 ± 0.5 | 4.62 ± 0.3 | 5.38 ± 0.4 | 6.65 ± 0.4 | 4.75 ± 0.3 |
| **Volume fraction (%)** | 6 ± 0.47 | 5.4 ± 0.84 | 4.4 ± 0.59 | 13.9 ± 0.7 | 27.1 ± 0.88 | 25.2 ± 4.64 |
| **Radius (µm)** | 1.4 ± 0.08 | 1.1 ± 0.04 | 1.1 ± 0.1 | 2.5 ± 0.04 | 3.2 ± 0.1 | 3.5 ± 0.26 |
| **Connectivity** | 0.84 ± 0.033 | 0.85 ± 0.023 | 0.88 ± 0.008 | 0.93 ± 0.033 | 0.99 ± 9.5e-5 | 0.99 ± 0.003 |
| **Branch length (µm)** | 8 ± 1.4 | 8.1 ± 0.9 | 8.3 ± 0.5 | 12.8 ± 0.1 | 12.8 ± 0.2 | 14.9 ± 0.6 |
| **Junction density (number/µm)** | 0.06 ± 0.001 | 0.06 ± 0.001 | 0.06 ± 0.002 | 0.03 ± 0.001 | 0.04 ± 9.4e-5 | 0.04 ± 0.001 |

**Supp table 02: Morphometric characterization of hepatocytes from early postnatal development to adulthood.**

HEPATOCYTES (GENERAL)

| | P1 | P16 | Adult |
|---|---|---|---|
| **Total number/mm³** | 474336 ± 35903 | 595904 ± 33765 | 235756 ± 31938 |
| **Volume fraction (%)** | 58.5 ± 1.4 | 59.6 ± 0.56 | 56.7 ± 2.22 |
| **Cell volume (µm³)** | 2152 ± 160 | 1731 ± 94 | 4471 ± 525 |
| **Mono-nuclear (%)** | 75.8 ± 4.1 | 79.3 ± 7.6 | 63.1 ± 7.5 |
| **Bi-nuclear (%)** | 17.3 ± 2.1 | 16.9 ± 4.7 | 34.5 ± 7.1 |
| **Nucleus volume (µm³)** | 344 ± 47 | 342 ± 40 | 480 ± 71 |
| **Nucleus elongation** | 0.83 ± 0.050 | 0.52 ± 0.061 | 0.4 ± 0.057 |

HEPATOCYTES (BY PLOIDY)

| | **1 x 2n** | | |
|---|---|---|---|
| | **P1** | **P16** | **Adult** |
| **Percentage of total hepatocytes (%)** | 58.9 ± 2.7 | 52.4 ± 1.2 | 27.4 ± 0.8 |
| **Cell volume (μm³)** | 1412 ± 82 | 1247 ± 128 | 2175 ± 291 |
| **Cell elongation** | 0.78 ± 0.006 | 0.59 ± 0.025 | 0.65 ± 0.015 |
| **Apical surface (%)** | 15.4 ± 0.5 | 18.2 ± 0.8 | 17.8 ± 3.3 |
| **Basal surface (%)** | 23.1 ± 1.2 | 39.1 ± 2.7 | 32.5 ± 2.7 |
| **Lateral surface (%)** | 61.5 ± 1.4 | 42.7 ± 3.6 | 49.7 ± 5.9 |
| **Number of neighbours** | 9 ± 0.24 | 8 ± 0.48 | 8 ± 0.33 |

| | **1 x 4n** | | |
|---|---|---|---|
| | **P1** | **P16** | **Adult** |
| **Percentage of total hepatocytes (%)** | 15.5 ± 4.3 | 24.9 ± 5.5 | 27.9 ± 4.4 |
| **Cell volume (μm³)** | 1821 ± 329 | 1612 ± 107 | 3756 ± 410 |
| **Cell elongation** | 0.94 ± 0.022 | 0.59 ± 0.026 | 0.57 ± 0.054 |
| **Apical surface (%)** | 15.8 ± 0.9 | 18.9 ± 0.5 | 19.8 ± 3.0 |
| **Basal surface (%)** | 23.3 ± 2.5 | 40.6 ± 3.1 | 39.6 ± 4 |
| **Lateral surface (%)** | 60.9 ± 3.4 | 40.5 ± 3.6 | 40.6 ± 7 |
| **Number of neighbours** | 10 ± 0.69 | 8 ± 0.57 | 9 ± 0.57 |

| | **1 x 8n** | | |
|---|---|---|---|
| | **P1** | **P16** | **Adult** |
| **Percentage of total hepatocytes (%)** | 1.4 ± 1.2 | 1.6 ± 0.8 | 7.1 ± 3.7 |
| **Cell volume (μm³)** | 2801 ± 286 | 3250 ± 483 | 5949 ± 551 |
| **Cell elongation** | 1.06 ± 0.141 | 0.75 ± 0.065 | 0.52 ± 0.050 |
| **Apical surface (%)** | 16.9 ± 3 | 13.2 ± 2.5 | 22.7 ± 2.3 |
| **Basal surface (%)** | 23.8 ± 2.9 | 50.2 ± 7.1 | 42.8 ± 3.8 |
| **Lateral surface (%)** | 59.3 ± 5.8 | 36.6 ± 6.7 | 34.5 ± 6.1 |
| **Number of neighbours** | 11 ± 1.4 | 11 ± 0.33 | 11 ± 0.93 |

| | **2 x 2n** | | |
|---|---|---|---|
| | **P1** | **P16** | **Adult** |
| **Percentage of total hepatocytes (%)** | 13.6 ± 2.7 | 12.6 ± 5.1 | 15.0 ± 2.8 |
| **Cell volume (µm³)** | 3352 ± 178 | 2586 ± 308 | 4474 ± 529 |
| **Cell elongation** | 1.07 ± 0.038 | 0.93 ± 0.030 | 0.65 ± 0.050 |
| **Apical surface (%)** | 19.5 ± 1.7 | 19.7 ± 1.8 | 23.3 ± 3 |
| **Basal surface (%)** | 25.6 ± 0.9 | 41.1 ± 2.2 | 38.0 ± 3.6 |
| **Lateral surface (%)** | 54.9 ± 2.6 | 39.2 ± 3.9 | 38.7 ± 6.3 |
| **Number of neighbours** | 13 ± 0.02 | 10 ± 0.45 | 11 ± 0.68 |

| | **2 x 4n** | | |
|---|---|---|---|
| | **P1** | **P16** | **Adult** |
| **Percentage of total hepatocytes (%)** | 0.5 ± 0.2 | 1.6 ± 0.6 | 15.7 ± 4.5 |
| **Cell volume (µm³)** | 3533 ± 782 | 3220 ± 349 | 7279 ± 596 |
| **Cell elongation** | 1.06 ± 0.112 | 1.12 ± 0.038 | 0.67 ± 0.058 |
| **Apical surface (%)** | 19.3 ± 1.2 | 21.3 ± 1.3 | 22.3 ± 2.1 |
| **Basal surface (%)** | 21.9 ± 8.6 | 43.5 ± 2.9 | 44.7 ± 4.7 |
| **Lateral surface (%)** | 58.7 ± 7.5 | 35.3 ± 4.2 | 33 ± 6.6 |
| **Number of neighbours** | 13 ± 1.22 | 12 ± 0.97 | 12 ± 0.49 |

| | **2 x 8n** | | |
|---|---|---|---|
| | **P1** | **P16** | **Adult** |
| **Percentage of total hepatocytes (%)** | 0 ± 0 | 0.1 ± 0.7 | 1.0 ± 0.5 |
| **Cell volume (µm³)** | 0 ± 0 | 1482 ± 1482 | 9792 ± 603 |
| **Cell elongation** | 0 ± 0 | 0.38 ± 0.377 | 0.84 ± 0.144 |
| **Apical surface (%)** | 0 ± 0 | 9 ± 9 | 17.4 ± 3.7 |
| **Basal surface (%)** | 0 ± 0 | 11.5 ± 11.5 | 48.6 ± 7.4 |
| **Lateral surface (%)** | 0 ± 0 | 12.9 ± 12.9 | 34.0 ± 10.3 |
| **Number of neighbours** | 0 ± 0 | 4 ± 4 | 13 ± 0.5 |

**Supp table 03: Morphometric characterization of non-parenchymal cells from early postnatal development to adulthood.**

| | HSC | | | KC | | |
|---|---|---|---|---|---|---|
| | **P1** | **P16** | **Adult** | **P1** | **P16** | **Adult** |
| **Total number/mm$^3$** | 61449 ± 11523 | 63983 ± 4805 | 44217 ± 7397 | 107528 ± 17238 | 29785 ± 5343 | 18986 ± 3851 |
| **Volume fraction (%)** | 9.2 ± 1.87 | 9.8 ± 1.84 | 7.3 ± 0.49 | 8.8 ± 1.93 | 7 ± 1.15 | 3.6 ± 0.04 |
| **Cell volume (µm$^3$)** | 1172 ± 64 | 1303 ± 274 | 1578 ± 95 | 2170 ± 386 | 1934 ± 246 | 1702 ± 86 |
| **Cell elongation** | 2.73 ± 0.006 | 2.52 ± 0.066 | 2.51 ± 0.118 | 1.7 ± 0.127 | 2.07 ± 0.120 | 2.34 ± 0.045 |
| **Nucleus volume (µm$^3$)** | 325 ± 31 | 262 ± 45 | 193 ± 52 | 352 ± 70 | 260 ± 42 | 151 ± 36 |
| **Nucleus elongation** | 1.1 ± 0.03 | 1.18 ± 0.1 | 1.02 ± 0.12 | 1.01 ± 0.09 | 1.36 ± 0.21 | 1.41 ± 0.3 |

### Supp table 04: Morphometric analysis of BC and sinusoidal networks in absence of KCs

| | Bile canaliculi | | Sinusoids | |
|---|---|---|---|---|
| | **PBS** | **Clodronate** | **PBS** | **Clodronate** |
| **Total length (m/mm³)** | 4.94 ± 0.1 | 5.37 ± 0.2 | 4.41 ± 0.3 | 3.87 ± 0.8 |
| **Volume fraction (%)** | 6.9 ± 0.72 | 5.66 ± 0.55 | 22 ± 1.84 | 23.2 ± 4.76 |
| **Radius (µm)** | 1.4 ± 0.11 | 1.2 ± 0.13 | 3.3 ± 0.16 | 3.7 ± 0.69 |
| **Connectivity** | 0.88 ± 0.018 | 0.91 ± 0.012 | 0.97 ± 0.012 | 0.96 ± 0.029 |
| **Branch length (µm)** | 8.3 ± 0.8 | 8.0 ± 0.3 | 14.3 ± 0.1 | 13.6 ± 0.7 |
| **Junction density (number/µm)** | 0.06 ± 0.001 | 0.06 ± 0.002 | 0.03 ± 0.001 | 0.03 ± 0.004 |

### Supp table 05: Morphometric characterization of hepatocytes in absence of KC

HEPATOCYTES (GENERAL)

| | **PBS** | **Clodronate** |
|---|---|---|
| **Total number/mm³** | 302397 ± 26303 | 318449 ± 44948 |
| **Volume fraction (%)** | 57.4 ± 1.86 | 57.7 ± 2.48 |
| **Cell volume (µm³)** | 3224 ± 260 | 3321 ± 461 |
| **Mono-nuclear (%)** | 69.6 ± 0.8 | 51.4 ± 4.9 |
| **Bi-nuclear (%)** | 24.3 ± 1.5 | 39.3 ± 4.9 |
| **Nucleus volume (µm³)** | 433 ± 58 | 436 ± 36 |
| **Nucleus elongation** | 0.62 ± 0.068 | 0.49 ± 0.049 |

HEPATOCYTES (BY PLOIDY)

| | 1 x 2n | |
|---|---|---|
| | **PBS** | **Clodronate** |
| **Percentage of total hepatocytes (%)** | 49.9 ± 3.8 | 31.9 ± 6.3 |
| **Cell volume (µm³)** | 1922 ± 86 | 1690 ± 172 |
| **Cell elongation** | 0.73 ± 0.025 | 0.71 ± 0.035 |
| **Apical surface (%)** | 21.3 ± 3.1 | 17.4 ± 1.7 |
| **Basal surface (%)** | 35.1 ± 2.5 | 37.1 ± 4.1 |
| **Lateral surface (%)** | 43.6 ± 5 | 45.5 ± 3.9 |
| **Number of neighbours** | 8 ± 0.48 | 8 ± 0.78 |

| | 1 x 4n | |
|---|---|---|
| | **PBS** | **Clodronate** |
| **Percentage of total hepatocytes (%)** | 18.5 ± 2.8 | 17.1 ± 3.3 |
| **Cell volume (µm³)** | 3123 ± 143 | 2517 ± 161 |
| **Cell elongation** | 0.68 ± 0.013 | 0.68 ± 0.066 |
| **Apical surface (%)** | 24.5 ± 2.2 | 21.9 ± 1.8 |
| **Basal surface (%)** | 38.7 ± 3.2 | 38.4 ± 6.1 |
| **Lateral surface (%)** | 36.9 ± 4.6 | 39.7 ± 7.3 |
| **Number of neighbours** | 9 ± 0.68 | 9 ± 0.93 |

| | 1 x 8n | |
|---|---|---|
| | **PBS** | **Clodronate** |
| **Percentage of total hepatocytes (%)** | 1.1 ± 0.2 | 2.1 ± 1.4 |
| **Cell volume (µm³)** | 3972 ± 329 | 4388 ± 530 |
| **Cell elongation** | 0.57 ± 0.038 | 0.63 ± 0.047 |
| **Apical surface (%)** | 28.4 ± 1.7 | 26.5 ± 1.3 |
| **Basal surface (%)** | 35.6 ± 2.7 | 31.0 ± 8.5 |
| **Lateral surface (%)** | 36.0 ± 2.0 | 42.5 ± 8.9 |
| **Number of neighbours** | 11 ± 0.48 | 11 ± 1.04 |

| | 2 x 2n | |
|---|---|---|
| | **PBS** | **Clodronate** |
| **Percentage of total hepatocytes (%)** | 15.3 ± 1.9 | 22.9 ± 3.6 |
| **Cell volume (µm³)** | 4048 ± 126 | 3296 ± 354 |
| **Cell elongation** | 0.8 ± 0.003 | 0.82 ± 0.054 |
| **Apical surface (%)** | 25.9 ± 2.5 | 21.9 ± 0.9 |
| **Basal surface (%)** | 39.7 ± 2.8 | 42 ± 5.9 |
| **Lateral surface (%)** | 34.4 ± 4.8 | 36.2 ± 6.1 |
| **Number of neighbours** | 11 ± 0.84 | 10 ± 1.07 |

| | 2 x 4n | |
|---|---|---|
| | **PBS** | **Clodronate** |
| **Percentage of total hepatocytes (%)** | 5.4 ± 2.1 | 11.0 ± 3.1 |
| **Cell volume (µm³)** | 6417 ± 504 | 5289 ± 817 |
| **Cell elongation** | 0.78 ± 0.056 | 0.79 ± 0.033 |
| **Apical surface (%)** | 30.5 ± 0.8 | 24.2 ± 1 |
| **Basal surface (%)** | 41.0 ± 5 | 40.9 ± 7.3 |
| **Lateral surface (%)** | 28.5 ± 4.2 | 34.9 ± 6.8 |
| **Number of neighbours** | 13 ± 1.07 | 12 ± 1.32 |

| | 2 x 8n | |
|---|---|---|
| | **PBS** | **Clodronate** |
| **Percentage of total hepatocytes (%)** | 0.4 ± 0.2 | 0.7 ± 0.2 |
| **Cell volume (µm³)** | 7471 ± 1438 | 5175 ± 1918 |
| **Cell elongation** | 0.67 ± 0.082 | 0.88 ± 0.374 |
| **Apical surface (%)** | 26.4 ± 2.6 | 25.1 ± 4.2 |
| **Basal surface (%)** | 41.3 ± 1.5 | 39.9 ± 2.6 |
| **Lateral surface (%)** | 32.3 ± 3.6 | 35.1 ± 6.7 |
| **Number of neighbours** | 16 ± 1.11 | 12 ± 2.34 |

### Supp table 06: Morphometric characterization of HSCs in absence of KCs

| | HSC | | KC | |
|---|---|---|---|---|
| | **PBS** | **Clodronate** | **PBS** | **Clodronate** |
| **Total number/mm³** | 40851 ± 11114 | 90995 ± 24403 | 23657 ± 1854 | - |
| **Volume fraction (%)** | 6.5 ± 1.29 | 9.4 ± 2.63 | 4.8 ± 0.37 | - |
| **Cell volume (µm³)** | 1334 ± 117 | 1263 ± 135 | 1753 ± 290 | - |
| **Cell elongation** | 3.02 ± 0.191 | 2.49 ± 0.147 | 2.44 ± 0.079 | - |
| **Nucleus volume (µm³)** | 307 ± 35 | 308 ± 41 | 323 ± 35 | - |
| **Nucleus elongation** | 1.2 ± 0.097 | 0.95 ± 0.100 | 1.2 ± 0.046 | - |

### Supp table 07: Antibodies, dyes and reagents used for staining and experiments

| Reagent type (species) or resource | Designation | Source or reference | Identifiers | Dilution or concentration | Description |
|---|---|---|---|---|---|
| Primary antibodies | anti-Flk1 (goat polyclonal) | R&D System | AF644 | (1:100) | Sinusoidal marker |
| | anti-CD13 (rat monoclonal) | Novus | NB100-64843 | (1:500) | Canaliculi marker |
| | anti-F4/80 (rat monoclonal) | Abcam | ab6640 | (1:400) | F4/80$^{+}$ cells marker |
| | anti-Desmin (rabbit polyclonal) | Abcam | ab15200 | (1:200) | Stellate cells marker |
| Secondary antibodies | Donkey anti-Goat Alexa Fluor 647 | Invitrogen | A-21447 | (1:1000) | |
| | Donkey anti-Rat CF 568 | Biotium | 20092 | (1:1000) | |
| | Donkey anti-Rabbit Alexa Fluor Plus 647 | Invitrogen | A32795 | (1:1000) | |
| Small dyes | Phalloidin Alexa Fluor 647 | Invitrogen | A22287 | (1:1000) | Cell border |
| | Phalloidin Alexa Fluor 488 | Invitrogen | A12379 | (1:100) | Cell border |
| | DAPI | Invitrogen | D1306 | (1:1000) | Nuclei |
| Reagent | Clodronate and control (PBS) liposome | LIPOSOMA | CP-010-010 | (5 mg/ml) | |
